# Substrate-dependent oligomerization modulates DGAT1 activity and subcellular localization

**DOI:** 10.64898/2026.03.31.715571

**Authors:** Jennifer Sapia, Veijo T. Salo, Atiya Tahira Tasnim, Pablo Campomanes, Xuewu Sui, Stefano Vanni

**Affiliations:** Department of Biology, University of Fribourg, Chemin du Musée 10, 1700 Fribourg, Switzerland; Molecular Systems Biology Unit, European Molecular Biology Laboratory (EMBL), Heidelberg, Germany; Department of Biochemistry and Biophysics, College of Agriculture and Life Sciences, Texas A&M University, College Station, TX, USA; Cell Biology and Biophysics Unit, EMBL, Heidelberg, Germany; National Center of Competence in Research Bio-inspired Materials, University of Fribourg, 1700 Fribourg, Switzerland

**Keywords:** Lipid droplets, triglycerides, diacylglycerol, membrane curvature, lipid storage, diacylglycerol O-acyltransferase 1, DGAT1, molecular dynamics

## Abstract

Lipid Droplets (LDs) are ubiquitous organelles that are responsible for intracellular energy storage, in the form of highly esterified lipids such as triglycerides (TGs) and sterol esters. LDs emerge from and engage in stable contact sites with the endoplasmic reticulum (ER), where TG biosynthesis takes place by the action of the acyltransferase DGAT1. Despite the recent cryo-EM determination of the human dimeric structure of DGAT1^1,2^, many aspects of the mechanism underlying TG synthesis in the ER remain unclear. Using a combination of molecular dynamics (MD) simulations, biochemical reconstitutions and fluorescence microscopy in live cells, we characterize several steps of DGAT1 molecular mechanism. We found that DAG preferentially enters the catalytic pocket of DGAT1 from the ER luminal leaflet, via a pathway that involves several conserved residues. Each DGAT1 subunit is able to bind multiple DAG molecules, and the presence of DAG promotes the formation of high-order DGAT1 oligomers. DGAT1 displays a preference for curved bilayers *in silico*, and it preferentially localizes in the ER tubular network, where LD formation is proposed to take place, upon the increase in its natural substrate diacylglycerol (DAG). Overall, our investigations provide a molecular view of how the interplay between protein oligomerization, subcellular localization and substrate biophysical properties modulate DGAT1 enzymatic activity.

## Introduction

Lipids are essential building blocks of the cell, and they constitute an important energy source due to their high caloric density^3^. Excess lipids are stored in their esterified, neutral, form, as triglycerides (TGs) and sterol esters (SEs) in special structures called lipid droplets (LDs). LDs are intracellular organelles with a unique architecture, consisting of a neutral lipid (NL) core surrounded by a phospholipid monolayer enriched with a variety of LD-specific proteins^4–7^. As such, they are responsible for fat storage in the cell, acting as a buffer against the cytotoxicity of various compounds such as free fatty acids or oxidized lipids^5–8^, and as energy reservoir that can be later accessed to perform basal functions when food intake is reduced or insufficient^6^. NLs can be later catabolized by lipases to promote the incorporation of free fatty acids into phospholipids (PLs), and hence membrane synthesis and expansion. Because of this central role in lipid metabolism, LDs are involved in numerous diseases, such as metabolic diseases^9^, cancer^10^, viral infections^11^, diabetes and liver diseases^12–14^.

In mammalian cells, NL synthesis occurs across multiple cellular sites. In the endoplasmic reticulum (ER), TG synthesis is catalyzed by DGAT1 (acyl-coenzyme A: diacylglycerol O-acyltransferase 1) in the final step of two major TG biosynthetic pathways: the *de novo* glycerolipid synthesis pathway, also known as the Kennedy pathway, and the monoacylglycerol pathway^6^. DGAT1, a member of the membrane-bound O-acyltransferase (MBOAT) enzyme family^15^, is responsible of the transfer of an acyl group from acyl-CoA molecule to diacylglycerol (DAG), its most abundant natural lipid substrate. In addition to DGAT1, DGAT2, a second acyltransferase which is encoded by a gene of different family^16^, contributes to TG synthesis, functioning predominantly at the LDs^17,18^.

A detailed understanding of how TG are synthesized in the ER, however, has remained elusive for a long time: despite being cloned for the first time more than 20 years ago^19^, many aspects of DGAT1 mechanism remain unclear. The 3D structure of DGAT1 has been recently solved using cryogenic electron microscopy (cryo-EM)^1,2^, leading to a better understanding of the overall protein architecture and to the identification of the binding pocket for one of its two substrates, a fatty acyl-CoA ligand. On the other hand, the localization of its second substrate, DAG, as well as entry and exit pathways for both substrates and products remain uncharacterized. Finally, the interplay between DGAT1 activity, oligomerization state and ER membrane properties, as well as its consequences on LD formation, have not been investigated so far. All these uncertainties regarding to its molecular mechanism hinder the design of suitable drugs to modulate its mechanism of action.

In this study, we overcome these limitations by employing a combination of *in silico* approaches, complemented by *in vitro* and *in vivo* experiments. All-Atom (AA) molecular dynamics (MD) simulations of DGAT1 characterize the mechanism of DAG recruitment to the catalytic pocket and show that each DGAT1 monomer can bind multiple DAG molecules. Coarse-grained (CG) MD simulations identify a key role for DAG in modulating DGAT1 oligomerization state and membrane remodeling activity. *In vitro* reconstitution and crosslinking assays unveil that the presence of DAG promotes the formation of higher-order oligomers. *In vivo* light microscopy confirms the preference of DGAT1 for the ER tubular network and indicate that this preference is accentuated when DAG concentration increases. Taken together, our data suggest that the interplay between DGAT1, its substrate DAG, and the surrounding membrane environment create conditions that facilitate TG synthesis and, ultimately, LD biogenesis.

## Results

### DAG enters the DGAT1 catalytic pocket from the luminal leaflet

To gain a molecular understanding of DGAT1’s behavior in a native-like environment, we employed AA-MD simulations using the cryo-EM structure of the human DGAT1 dimer complexed with a molecule of oleoyl-Coenzyme A (CoA), one of its natural substrates, per each subunit^1^, as the starting model. We embedded the protein in an explicit lipid bilayer (Supplementary Fig. S1A), constituted by di-oleoyl phosphatidyl choline (DOPC) and di-oleoyl-glycerol (DAG), the latter being a second natural lipid substrate of DGAT1 and precursor of the final product TG. In all replicates, the dimeric structure of DGAT1 remains stable throughout the simulation while, within each DGAT1 pocket, the oleoyl-CoA molecule, and particularly its hydrophilic CoA moiety, appears displaced toward the cytosolic leaflet, potentially suggesting its possible entrance from the cytoplasm^20^ (Supplementary Fig. S1B).

In all simulations, we repeatedly observed that DAG molecules spontaneously enter the lateral tunnel within the transmembrane region of each DGAT1 monomer (Fig. 1A). We next investigated whether the substrate molecules follow a defined pathway toward the active site or whether its entry occurs in a more stochastic manner. Unlike a previous report observing one single DAG entering the DGAT1 pocket from the cytosolic leaflet^20^, in our multiple replicates we observe several instances of DAG entering DGAT1 pocket (Fig. 1B). The majority of them enter from the luminal side, while the cytosolic leaflet route was taken less frequently. (Fig. 1B).

**Figure 1.**
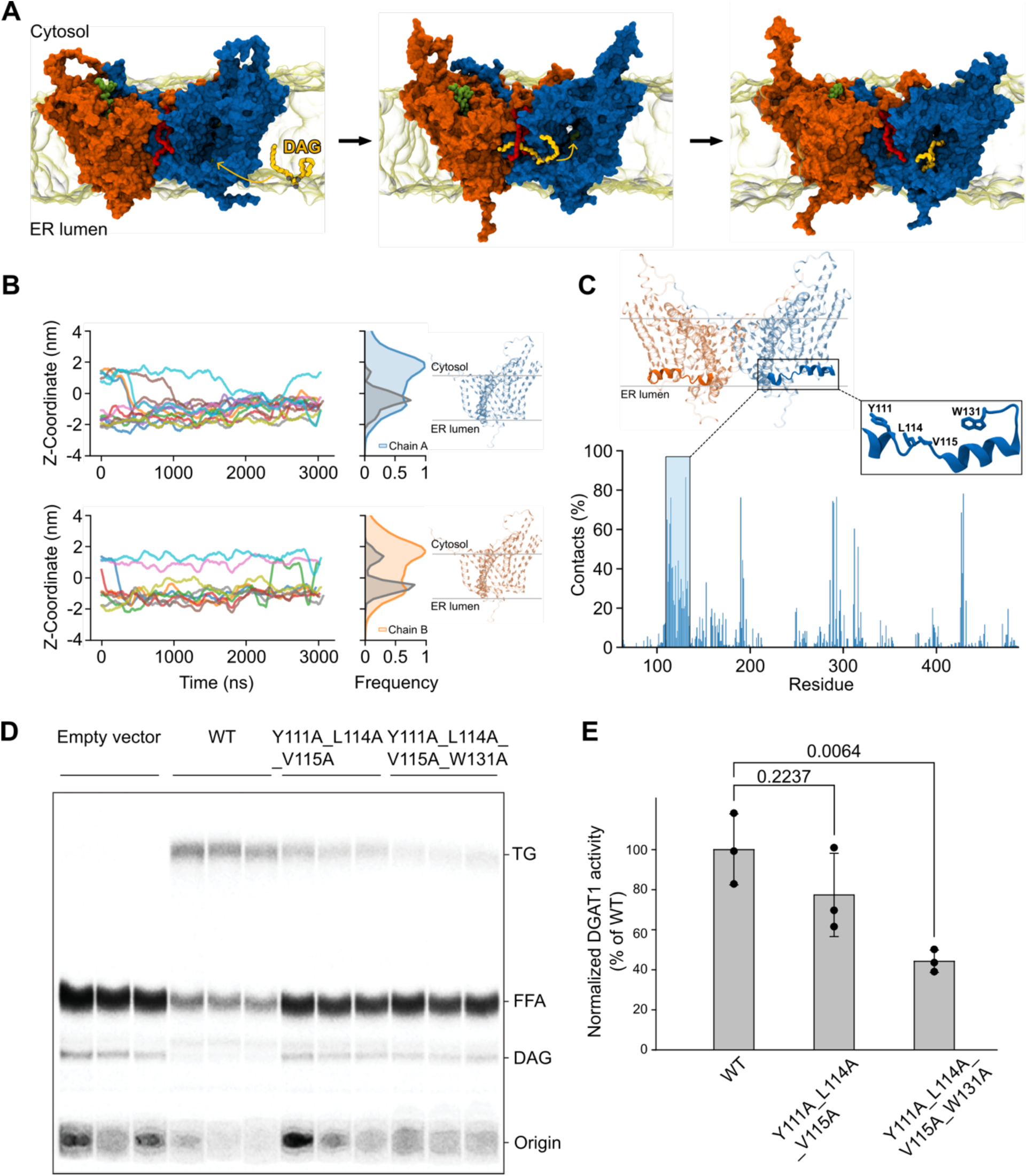
Characterization of DAG entry pathways into DGAT1 pocket: A) Schematic representation from AA simulations showing DAG (yellow) spontaneous entry within DGAT1 catalytic pocket. B) Time traces of DAG molecules z coordinates while entering the DGAT1 cavities along with density profiles for both bound DAG and DGAT1 monomers. Color code: left, each color represents the z coordinate of a single DAG molecule bound to DGAT1; right, the gray histogram representing the density of bound DAG overtime, while the blue and orange histogram represent respectively the DGAT1’s monomer A and B. C) Percentage of contacts between DAG molecules and each DGAT1’s residue. The inset shows the highly conserved residues, belonging to the ⍺-helix located at the bottom of the cavity, which interact more with the ligand. TLC analysis of TG formation by wide-type DGAT1 and mutations in the DAG entry pathway using yeast microsome overexpressing GFP-fused proteins. E) Quantification of the activity analyses shown in D after normalized with DGAT1 expression level (Fig S1C).

Analysis of DAG-protein contacts indicate that, during entry, DAG prominently interacts with several highly conserved residues, including Y111, L114, V115, and W131^1^, located on an amphipathic ⍺-helix at the base of the cavity (Fig. 1C). This result suggests a potential role of the ⍺-helix to regulate substrate entrance, and, therefore, DGAT1 catalytic activity. To test this hypothesis, we next used an established activity assay for DGAT1^1^. In short, using yeast microsome overexpressing human DGAT1 (Supplementary fig. S1C), we compared the activity of the wild-type with that of mutants in which the DAG-interacting hydrophobic residues were mutated into alanine.

These mutations significantly impair enzymatic activity, leading to an approximate 50% reduction compared to the WT (Fig. 1D-E), indicating that perturbations in the interaction between DAG molecules and the ⍺-helix along the entry pathway significantly decrease DGAT1 enzymatic activity.

### The catalytic pocket of DGAT1 can harbor multiple DAG molecules at the same time

We next investigate the position and localization of the DAG molecules inside the catalytic pocket. First, we observed that once bound inside DGAT1 cavity, in some instances DAG positions itself in very close proximity to the second natural substrate, oleoyl-CoA (Fig. 2A), resembling the lipid-like EM density observed in the cryo-EM structure^1^. This allows the two substrates, DAG and acyl-CoA, to be positioned close to each other within the cavity not only facilitating their interaction during the catalytic process but also placing them near crucial residues implicated in catalysis, such as His415, Arg378 and Met434 (Fig.2A inset). The spatial arrangement observed in the simulations underscores a structural framework that may be essential for the enzymatic function of DGAT1.

**Figure 2.**
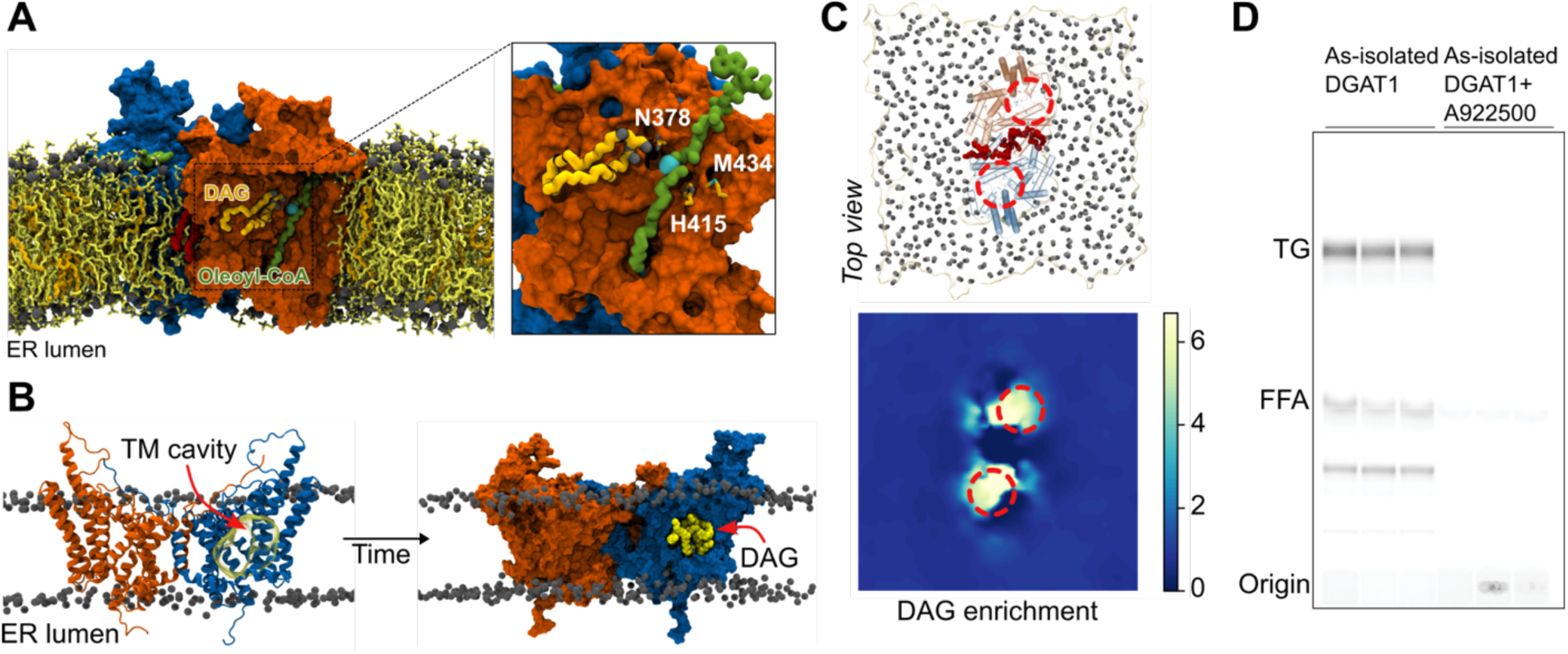
DAG recruitment into DGAT1 catalytic pocket: A) Representative snapshot from AA-MD simulations showing a DAG molecule (light orange, licorice representation) in the next proximity near oleoyl-CoA (green, Van der Waals representation). The inset shows detail from AA-MD simulations displaying the proximity of the two substrate, DAG and oleoyl-CoA, to several key catalytic residues. B) DGAT1 transmembrane cavity (left, yellow transparent surface) appears filled of DAG molecules (right, Vand der Waals) during the simulations. C) Top view (top) of DGAT1 embedded in a model bilayer and 2D enrichment map (bottom) showing accumulation of DAG inside DGAT1 cavities (red dashed circles). D) Purified DGAT1 is bound with endogenous DAG. Purified DGAT1 in GDN generated TG in the presence of oleoyl-CoA but without providing exogenous DAG substrate. The inhibitor treatment by A922500 completely abolished the TG production.

Next, we noticed that the large transmembrane cavity within each DGAT1 monomer is filled simultaneously with multiple DAG molecules, typically up to five (Fig. 2B-C). To test this unusual enzymatic configuration, we again performed DGAT1 activity assays, in the presence of only 14C-oleoyl CoA but not exogenous DAG (Fig. 2D, Supplementary fig. S1D). Of note, we observed substantial formation of 14C-TG, indicating that DAG molecules co-purified with DGAT1 (Fig. 2D). The presence of a DGAT1 inhibitor, on the other hand, completely blocks TG formation (Fig. 2D). All together these results confirm the presence of multiple DAG molecules inside the catalytic pocket that are readily available for catalysis.

### DGAT1 can induce and sense local curvature of its surrounding membrane environment

Since DAG can easily flip-flop between membrane leaflets^21–25^, the observed asymmetry between entry pathways (Fig. 1A) is intriguing, as the main asymmetry in the ER originates at the ER tubular network, where the two leaflets are intrinsically asymmetric due to the applied curvature stress^26,27^.

We noticed that the DGAT1 dimer has a pronounced conical shape, which is usually associated with a positive curvature of the bilayer^26,28^. Thus, we next investigated the positioning of the protein within the membrane and how it affects the surrounding environment. Our simulations indicate that the overall density of DGAT1 dimer appears mostly shifted towards the cytosolic leaflet, distributing asymmetrically within the lipid bilayer (Fig. 3A). Therefore, we next wondered whether the protein could impart some anisotropy to the membrane.

**Figure 3.**
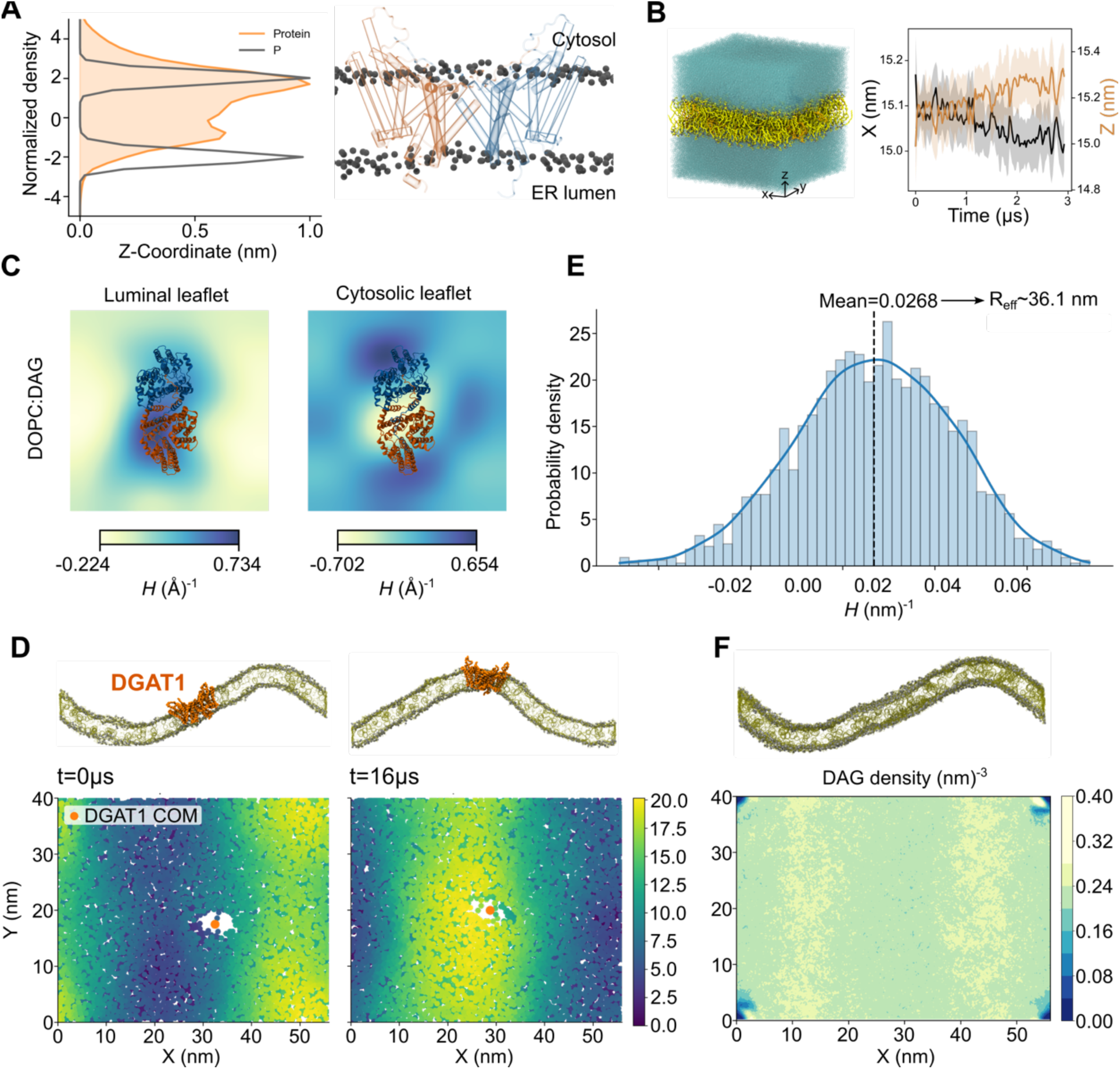
DGAT1 generates and sense membrane curvature: **A)** Density estimation of DGAT1 shows its localization within the bilayer. The protein is displaced towards the cytosolic leaflet of the membrane. **B)** Time evolution of the X (=Y) and Z dimensions of the simulated AA DGAT1-dimer embedded in a DOPC:DAG lipid bilayer, during the calculated trajectory. In the simulations, the Z dimension increases and the X dimension (as well as Y, which is not showed) decreases, indicating curvature generation. Each plot shows block-averages for consecutive 3 μs trajectory. **C)** Membrane curvature analysis of DOPC:DAG bilayer show membrane deformation, with the luminal leaflet adopting a negative curvature (yellow) and the cytosolic leaflet a positive curvature (blue). **D)** DGAT1 homodimer (orange) in a CG buckled bilayer (top panels) shows a propensity to localize in curved regions. The bottom panels show a top view of the DOPC phosphate beads, coloured based on their Z position (blue color corresponding to negative curvature, yellow to positive curvature). The center of mass (COM) of the DGAT1 dimer is shown as an orange dot. **E)** Histogram of mean curvature (*H*) indicates DGAT1 preference for positively highly-curved regions in a CG buckled membrane. The effective radius of curvature is also reported. **F)** Number density of DAG lipids showing their accumulations in curved regions of the buckled membrane (yellow areas).

To examine the impact of DGAT1 on its local environment, we next performed membrane curvature analysis which revealed that the DGAT1 dimer induces a local distortion in its immediate surroundings (Fig. 3B-C, Supplementary fig. S1E), generating significant curvature with a radius of approximately 20 nm, comparable to that observed in the ER tubular network^29^.

To further determine whether DGAT1 exhibits a preference for positively curved, flat, or negatively curved membrane regions^30,31^, we utilized CG-MD simulations of DGAT1 embedded in buckled bilayers. Again, the simulations revealed a clear preference for DGAT1 to localize in positively curved regions of the membrane (Fig. 3D-E, Supplementary fig. S1F-G) with an effective radius of curvature ranging around 37-40 nm, again in good agreement with reported experimental data^32^.

Furthermore, the same simulations showed that also DAG has a clear tendency to accumulate in the regions of the buckled membrane with high curvature (Fig. 3F, Supplementary fig. S1H), corroborating the idea that conical lipids preferentially partition in curved regions to reduce the membrane stress.

Taken together, our simulations unveil an interesting biophysical positive-feedback loop: due to their asymmetry, DGAT1 dimers display a preference for curved bilayers, in which their substrate, DAG, is naturally enriched (Fig. 3). There, DAG preferentially enters the catalytic pocket of DGAT1 from the negatively curved luminal leaflet (Fig. 1), in which it is naturally enriched due to its conical shape.

### DAG promotes DGAT1 localization to the ER tubular network

Based on the observed physical properties of DGAT1 and DAG, a preferential partitioning of DGAT1 to the ER tubular network could favor substrate access and, consequently, enzymatic activity. Furthermore, LD biogenesis has been shown to occur in curved regions of the ER membrane^33–35^, and coordination between TG synthesis by DGAT1 and LD formation at ER tubules could be advantageous for directing TG molecules inside LDs.

To directly test this hypothesis, we next co-transfected EGFP-tagged DGAT1 with the ER-luminal marker BFP-KDEL into human SUM159 breast cancer cells and analyzed its localization in ER sheets and tubules via live cell Airyscan microscopy. To do so, we used an analysis pipeline previously described^35^ to segment the ER network into sheets and tubules^35^ (Fig 4A). Relative to BFP-KDEL, DGAT1 displayed a slight enrichment at ER tubules at baseline and appeared to be relatively depleted from ER sheet regions (Fig 4B). This enrichment at tubules *vs.* sheets was significantly enhanced by treating cells with oleic acid to stimulate LD biogenesis (Fig 4B-C) and was further increased by co-treating the cells with DGAT inhibitors and oleic acid (Fig 4B-C), conditions in which DAG accumulates in the ER^36^. These data suggest that whilst in cells DGAT1 localization at baseline displays only slight preference for higher ER tubular regions, activating LD biogenesis and/or increasing DAG levels drive DGAT1 localization to ER tubular regions, harboring higher membrane curvature.

**Figure 4.**
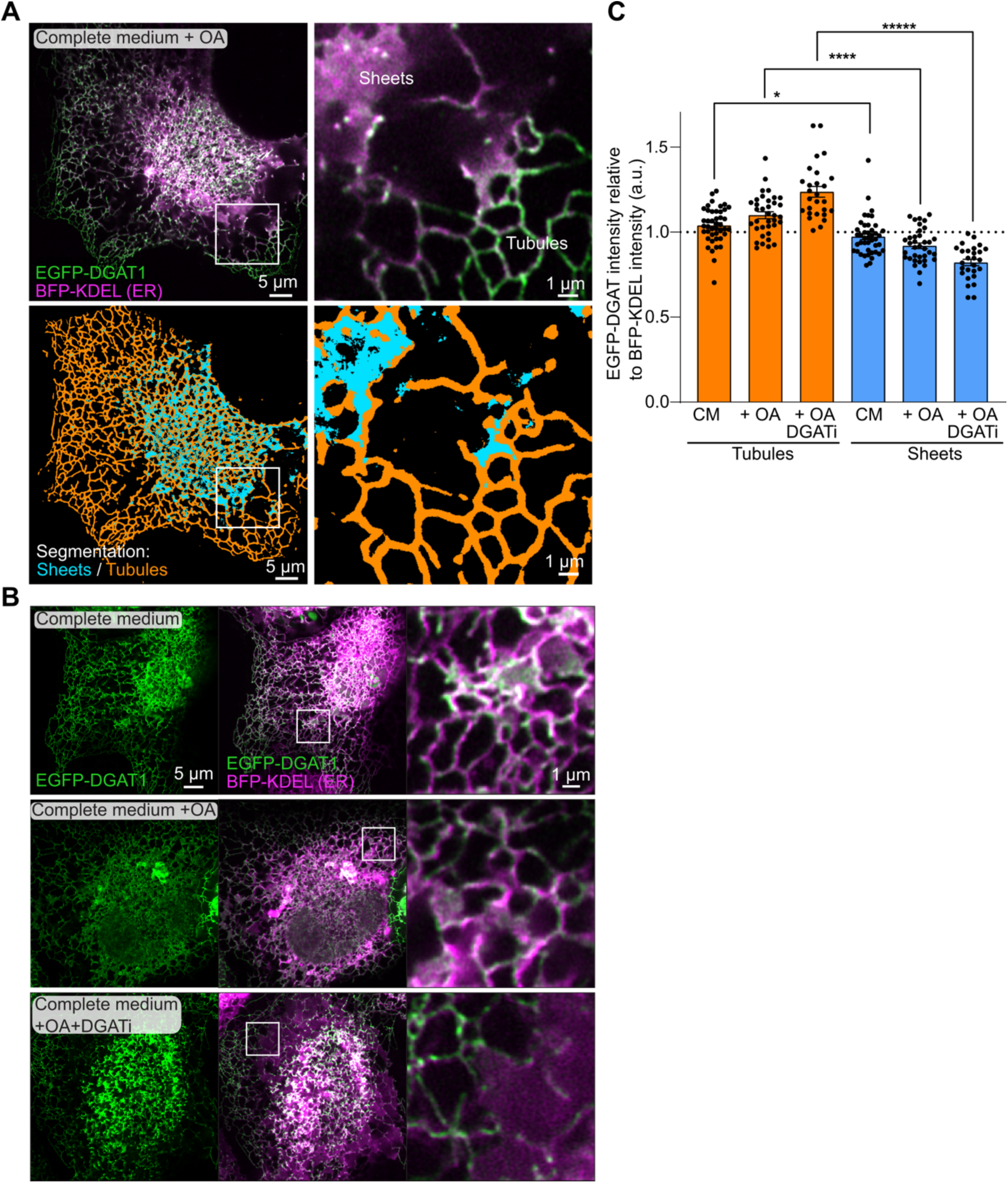
High DAG levels promote DGAT1 localization to ER tubules: **A)** SUM159 cells transfected with EGFP-DGAT1 and BFP-KDEL were treated with OA for 1 h and imaged live using Airyscan microscopy and the ER network was segmented into sheets and tubules. The EGPF-DGAT1 signal displays enrichment at peripheral tubular regions relative to sheet regions. **B)** Cells were treated and imaged as above in the conditions indicated in the figure. The enrichment of DGAT1 in tubules is more evident in oleic acid and oleic acid + DGAT inhibitor conditions. **C)** Analysis of (B). In the bar graph, each dot represents a cell. A value over 1 indicates EGFP-DGAT1 enrichment in the indicated ER compartment relative to BFP-KDEL, a value under 1 indicates de-enrichment. N= 24-40 cells per condition, 2 experiments. Statistics have been performed using Kruskal Wallis test followed by Dunn’s multiple comparison test, **** p < 0.0001. Of note, for each condition DGAT1 is significantly enriched at tubules vs sheets (* for complete medium, **** for complete medium and OA, **** for complete medium and OA and DGATi).

### DAG promotes DGAT1 oligomerization

Our data indicate that DGAT1 recruitment to the ER tubular network is driven by increasing DAG levels. To directly test this hypothesis and understand its underlying mechanistic details, we next performed CG-MD simulations in buckled bilayers utilizing multiple copies of DGAT1 dimer (Supplementary fig. S1I), in both the absence and presence of DAG at different concentrations (20%-30%) (Fig. 5A, Supplementary fig. S1J).

**Figure 5.**
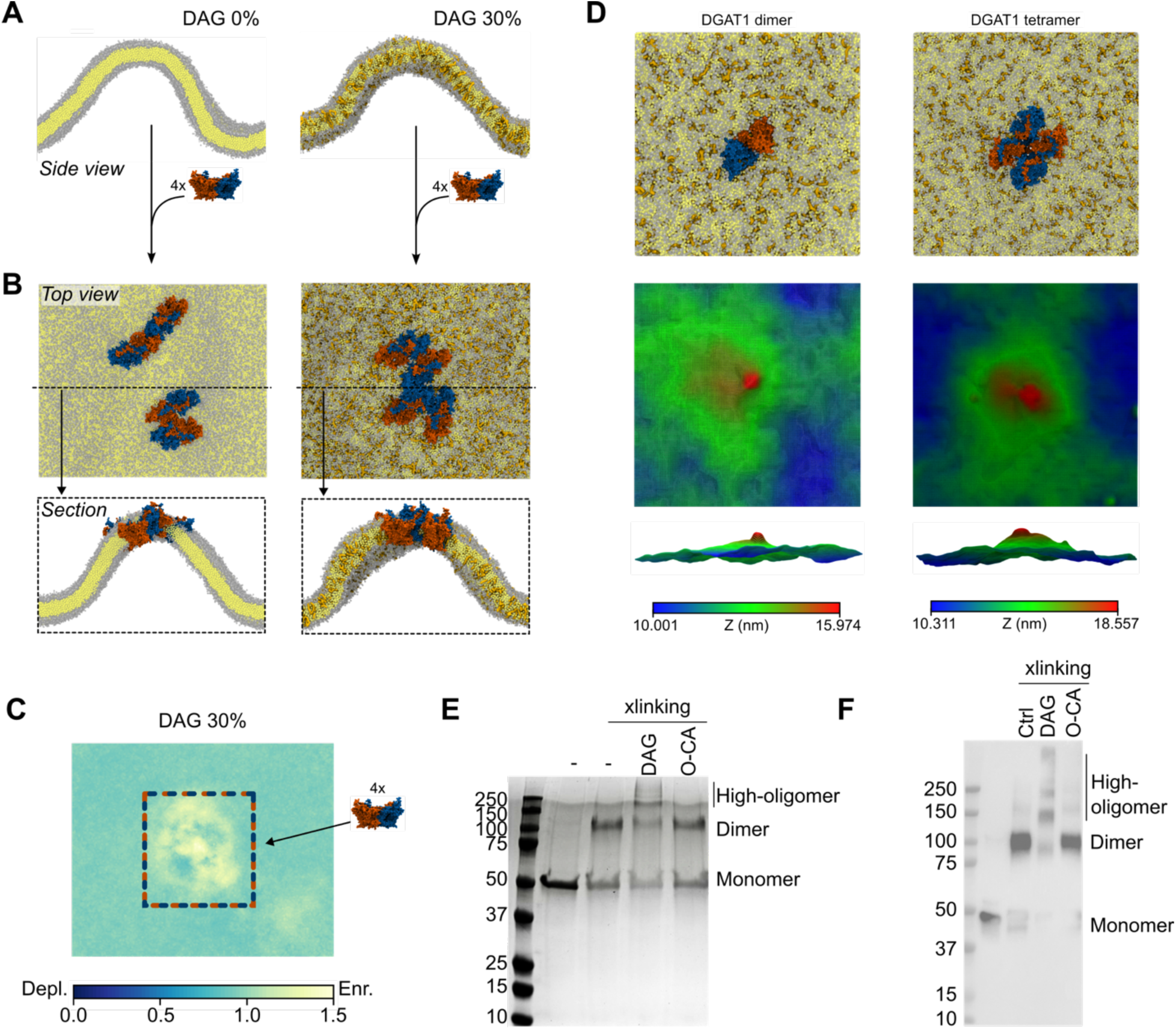
DAG promotes high-order DGAT1 oligomerization: **A)** Buckled membrane systems modelled in absence and in presence of DAG at 30% concentration. **B)** multiple copies of DGAT1 dimer clustering at the positively curved region. **C)** Enrichment map showing DAG accumulation in the proximity of the DGAT1 clusters 30% DAG concentration The dashed box represents the presence of DGAT1 dimers cluster at the region of DAG enrichment. The analysis is referred to one representative replica. **D)** Top view of DGAT1 dimer (left) and tetramer (right) embedded in a flat bilayer (upper panels) and corresponding surface grids (lower panels) showing membrane deformation caused by DGAT1 in different oligomeric states. **E)** DSS crosslinking and SDS-PAGE analyses of reconstituted DGAT1 after treating with DAG or oleoyl-CoA. **F)** WB analysis of crosslinking experiment using DGAT1 primary antibody. Different oligomerization states are labelled based on DGAT1 migration relative to the protein marker. The crosslinking assay were repeated three times independently with similar results.

Intringuingly, we observed that in the presence of high DAG concentrations, interactions between dimers led to the formation of high-order oligomers (Fig. 5B, Supplementary fig. S1J) that stably reside at the positively curved region of the buckled membrane.

In simulations performed in presence of DAG, the membrane surrounding these DGAT1 oligomers appear to be enriched with DAG molecules, indicating a possible role for the natural substrate in promoting DGAT1 oligomerization into high-order complexes (Fig. 5C, Supplementary fig. S1K). Further, by comparing the curvature preference of the DGAT1 dimer and tetramer in CG simulations (Fig. 5D, upper panels), we could observe that the tetramer displays enhanced membrane curvature preference (Fig. 5D, lower panels), suggesting that higher-order oligomers could have a higher preferential partitioning to highly curved ER tubules compared to DGAT1 dimers (Fig. 5B-C-D, Supplementary fig. S1I-J-K).

To directly test whether DAG can indeed promote the formation of high-order DGAT1 oligomers, we next incubated purified human DGAT1 with DAG and performed disuccinimidyl suberate (DSS) crosslinking experiments. SDS-PAGE analysis of purified DGAT1 predominantly migrated as dimer after DSS treatment, which was consistent with its dimeric structure as revealed by cryo-EM. However, upon preincubation with DAG, DGAT1 formed higher oligomers, and this oligomerization was specific to DAG, as oleoyl-CoA failed to induce higher oligomer formation (Fig 5E). To rule out that DAG-mediated higher oligomers formation was not simply due to DGAT1 aggregation, we performed western-blot (WB) analysis and found that DAG treatment indeed promoted DGAT1 oligomerization into different various oligomerization states in addition to dimer, including trimer, tetramer, and higher oligomers (Fig. 5F).

Collectively, these findings support a model in which the local accumulation of the DAG substrate within specific membrane regions (ER tubules) promotes the recruitment of the DGAT1 enzyme to those sites, where LD formation takes place, by modulating DGAT1 oligomeric state.

## Discussion

Despite its fundamental role in LD biosynthesis as a key enzyme in TG synthesis, DGAT1 has remained relatively understudied for many years. Only recently, with the advancements in cryo-EM, its 3D structure has been resolved. However, many uncertainties remain regarding its precise function within the broader context of LD biogenesis.

DGAT1 functions as a lipid remodeling enzyme, catalyzing the transfer of an acyl chain from an activated fatty acid, acyl-CoA, to a DAG molecule, to generate TG. DAG itself is a well-known lipid intermediate, playing a critical role in multiple metabolic pathways, including signal transduction cascades^37^. It is produced through lipid remodeling processes in the classical Kennedy pathway, either by the hydrolysis of phospholipids such as phosphatidic acid (PA) or through the *de novo* synthesis from its precursor, monoacylglycerol (MAG)^37^. More recently, an alternative, DGAT1/2 independent TG biosynthesis pathway involving a protein called DIESL and its regulator TMX1 has also been revealed^38^.

Beyond its role as a substrate in TG biosynthesis, DAG appears to influence the early stages of LD formation. Due to its inverted conical shape and its ability to spontaneously flip-flop between membrane leaflets^21–25^, DAG preferentially accumulates in membrane regions with high negative curvature, potentially also actively contributing to curvature generation. Previous studies suggest that DAG interacts with the FIT2 proteins in yeast, retaining DAG in the luminal leaflet and facilitating LD extrusion from the cytosolic leaflet^39^. In the absence of FIT2, LDs remain embedded in the ER^39^. Additionally, DAG-enriched ER regions recruit key proteins involved in LD biogenesis, such as Pex30^33^ and Nem1^40^, while MD simulations showed DAG clustering within human seipin rings^41,42^. In mammalian cells, DAG also promotes the recruitment of PLIN3, which associates with newly formed LDs emerging from the ER^43^.

In this study, we used computational and experimental approaches to reveal that DAG modulates DGAT1 activity not only as a substrate for the synthesis of TG, but also as a key regulator of its oligomerization state and intracellular localization. Our findings lead to a mechanistic model where DAG plays multiple pivotal roles to promote TG synthesis and LD biogenesis and growth. In detail, our data suggest that because of its biophysical properties, DAG molecules preferentially partition in the luminal leaflet of ER tubular regions. Similarly, DGAT1 also diffuses preferentially to ER tubules, as a consequence of the asymmetric protein shape of its most frequent dimeric conformation. Thanks to this coincident enrichment, DAG molecules can enter the DGAT1 catalytic cavity via a luminal pathway, where well-conserved residues localized at the luminal entrance of DGAT1 transmembrane pockets modulate substrate entrance. Enrichment of DAG molecules around DGAT1 also modulates protein oligomerization, promoting the formation of high-order oligomers. In turn, this further increases protein propensity for curved membrane regions, promoting a positive feedback loop that leads to further enrichment of DGAT1 in ER tubules.

Overall, since ER tubules are the main LD formation sites^35^, our data suggest that DAG-driven DGAT1 oligomerization and subcellular localization are likely to contribute, together with other proteins such as seipin, towards the spatial organization of LD formation within the cell. Our study unveils an important role for lipid biophysical properties in these processes, and it paves the way towards a better understanding of the early stages of LD formation and their associated regulatory mechanisms.

## Methods

### MD simulations

All MD simulations were performed using the cryo-EM structure of human DGAT1 dimer (pdb codes: 6VZ1^1^, 6VP0^1^), complexed with a molecule of oleoyl-CoA per each subunit. Structural lipids resolved along with the structure of the protein (e.g. four POPE molecules located at the dimer interface in 6VZ1) were kept during the simulations.

*AA simulations*. The DGAT1 dimer was embedded in a simple ER-like model membrane constituted of 85%DOPC-15%DAG. The systems were first generated using CHARMM-GUI membrane builder tool^44,45^, then minimized and equilibrated following the classical CHARMM-GUI 6 step-protocol. MD productions were carried out using GROMACS software^46^ and employing the CHARMM36m force-field^47–49^. Four replicates of 3 μs each were simulated, utilizing a timestep of 2 fs in the NPT ensemble. Temperature was maintained at 310 K using the Nosé-Hoover thermostat^50^, while the Parrinello-Rahman barostat^51^ was employed to keep the pressure at 1 bar and a compressibility of 4.5 x 10^-5^ bar^-1^ every 5 ps, applying a semi-isotropic pressure coupling. Long-range electrostatic interactions were treated using the Particle Mesh Ewald (PME) algorithm^52,53^, with a cutoff of 1.2 nm. Lennard-Jones (LJ) interactions were truncated at 1.2 nm. Linear Constraint Solver (LINCS) algorithm^54^ was applied to treat bond constraints.

The number of DAG molecules in each DGAT1 subunit and their position with respect to the z-axis were calculated employing a custom-built script. The frequency of interaction of DAG with each protein residue was defined by calculating the number of contacts amongst them overtime. The distance between the two natural ligands, DAG and oleoyl-CoA was monitored by using the GROMACS *gmx mindist* module.

DAG enrichment analysis was performed following the protocol described in ^55^: the 2D lateral density maps for each lipid species (DAG and DOPC) were first generated the *gmx densmap* module by GROMACS. The DAG enrichment was the calculated dividing DAG density by the sum of DAG and DOPC density and then again divided by the concentration of DAG in the bilayer.

The partial density analysis of diverse system components (e.g. protein, DOPC phosphates, oleoyl-CoA sulfur atoms) was calculated using *gmx density*.

2D maps of leaflets mean curvature were generated using the MDAnalysis MembraneCurvature tool (10.5281/ZENODO.5553452); the time evolution of the x,y,z dimensions of the AA-simulation box were performed utilizing *gmx energy* per each replica and then block-averaged.

*CG simulations.* The *apo* form of DGAT1 dimer was mapped to CG-MARTINI model using the *martinize2* tool^56^ and embedded in a pure DOPC buckled bilayer. To generate the buckled membrane, a rectangular shaped (50×30×30 nm), flat DOPC bilayer, solvated with MARTINI water and ionized with 150 mM of NaCl, was first generated using the *insane.py* script^57^. The system was first energy minimized employing the steepest descent algorithm and then equilibrated using NPT pressure coupling for 500 ns. To initiate the buckling, a lateral pressure of 2 bar was applied on the x direction, while keeping the y dimension fixed. This allows the system to expand only along the x and z directions. To do so, the anisotropic pressure coupling was employed, setting compressibility 𝑘x= 𝑘z=3×10^−5^ Pa, while 𝑘y= 0 Pa. Upon buckle generation, the curved bilayer was again equilibrated using semiisotropic conditions, with a pressure of 1 bar and the compressibility 𝑘x= 𝑘y= 0, 𝑘z = 3×10^−5^ Pa, allowing fluctuations only along the z dimension. MD production of two independent replicates of 16 μs each was run using GROMACS package, employing the MARTINI3 forcefield^58,59^. To maintain the pressure constant, the Parrinello-Rahman barostat^51^ was used, while the temperature was kept constant at 310 K employing the V-rescale thermostat^60^.

To investigate curvature-dependent DGAT1 oligomerization, CG-MD simulations were performed using a pre-equilibrated buckled membrane system in both the absence and presence (20% and 30%) of DAG molecules. An initial copy of DGAT1 dimer was manually positioned at the flat region corresponding to the transition zone between positive and negative curvature of the buckled bilayer. The system was then simulated until the enzyme spontaneously diffused towards the positively curved membrane region. Following this relocation, a second DGAT1 dimer unit was introduced at the same flat interface region, and the simulation was continued until it likewise diffused towards the positively curved area. This sequential insertion procedure was repeated for a total of four DGA1 dimer copies, each added only after the previously introduced enzyme had migrated away from the insertion site. Upon addition of the fourth enzyme, simulations (4 replicates per each membrane composition) were extended to ensure stabilization and convergence of the DGAT1 oligomer at the positively curved membrane region. The total accumulated simulation time from the insertion of the first enzyme to the convergence of the four-enzyme cluster was 50 μs per each replicate.

Simulations of both DGAT1 dimer and tetramer were run in presence of a flat bilayer composed by 85%DOPC-15%DAG. For DGAT1 dimer, the cryo-EM structure from ^1^ was utilized as starting point, while the DGAT1 tetrameric structure was predicted using Alphafold2^61,62^. Both the structures where first converted into CG-MARTINI model using *martinize2* and subsequently inserted in the flat membrane utilizing the *insane.py* script. The systems, solvated and ionized with 150 mM of NaCl, were minimized using the steepest descent algorithm and then shortly equilibrated for 10 ns, using semiisotropic conditions in the NPT ensemble and a timestep of 20 fs. 2 replicates for both systems containing dimer and tetramer were simulated for 10 µs each. Pressure was maintained constant utilizing the Parrinello-Rahman barostat, while the temperature was kept at 310K using the V-rescale thermostat.

Mean curvature analysis for the buckled membranes was performed using the MemCurv Python package^31^, while the DAG number density was calculated utilizing the GROMACS module *gmx densmap*. Membrane deformation was estimated using the tool Suave^63^.

All images deriving from MD simulations were rendered using Visual Molecular Dynamics (VMD) software^64^, while all plots were produced using the python module *matplotlib*^65^.

### In vivo assays

Sum159 cells were maintained in DMEM/F-12 GlutaMAX (Thermo Fisher Scientific) supplemented with 5 mg ml^−1^ insulin (Cell Applications), 1 mg ml^−1^ hydrocortisone (Sigma), 5% FBS (v/v), 50 mg ml^−1^ streptomycin and 50 U ml−1 penicillin. All cells were cultured at 37 °C in a 5% CO2-containing atmosphere. These conditions were used for all live cell imaging experiments. 10 mM stock oleic acid was prepared in PBS and used at a final concentration of 500 µM. DGAT1 (PZ0207) and DGAT2 (PZ0233) inhibitors were from Sigma and used at a final concentration of 5 µM.

For experiments, cells were seeded onto Ibidi 8-well dishes (15 000 cells per well) and transfected 1 day later with EGFP-DGAT1 (gift from Tobias Walther and Robert Farese) and BFP-KDEL (gift from Elina Ikonen, Addgene 87163^66^) using Lipofectamine LTX with Plus Reagent according to manufacturer instructions. 1 day later, live cell imaging was performed in Fluorobrite DMEM medium supplemented as above. For oleic acid and DGAT inhibitor treatments, cells were treated with oleic acid and the inhibitors for 45-90 min during the imaging session. Single z-plane images on the ER plane between the nucleus and the coverslip, covering the whole cell, were acquired on a Zeiss LSM880 confocal microscope, with an Airyscan detector and Plan-Apochromat ×63/1.4 numerical aperture (NA) Oil DIC M27 objective. Images were processed with identical settings in each data set using Zen Blue (Zeiss).

Analysis of ER morphology was done as described^35^. Briefly, extracellular background levels for each channel were acquired my measuring the intensity in multiple regions outside cells and background subtracted intensities were used for subsequent analyses. Single cells were cropped manually using ImageJ FIJI. Then, the BFP-KDEL signal was segmented into background, ER tubules, ER sheets and nuclear envelope using ilastik^67^ in pixel classification mode. For each cell, the mean background subtracted intensity of EGFP-DGAT1 in either sheets or tubules was compared to the mean background subtracted intensity of BFP-KDEL in either sheets or tubules in that cell to obtain the relative enrichment ratio in either sheets or tubules, using CellProfiler and custom-made MATLAB scripts described in ^68,69^.This method has been shown to recapitulate the known enrichment of well-characterised ER maker proteins (REEP-5, CLIMP-63, SEC61b) in sheets or tubules.

### DGAT1 overexpression and crosslinking

For DGAT1 overexpression in yeast, codon optimized human *DGAT1* gene was cloned into pDDGFP_LEU2 vectors (Addgene plasmid no. 102334) harboring a modified EGFP tag at the N-terminus of DGAT1. All the DGAT1 mutants were generated by the QhickChange mutagenesis kit (Agilent) using the protocol provided by the manufacturer. The plasmids were transformed into *S. cerevisiae* strain BJ5460 (ATCC no. 208285, genotype: *MATa ura3-52 trp1 lys2-801 leu2-delta1 his3-delta200 pep4::HIS3 prb1-delta1.6R can1 GAL*) using the lithium acetate method^70^. Transformants grew for ∼48 h on plates with yeast nitrogen base (YNB), complete supplement mixture (CSM) lacking uracil, and 2% w/v glucose before streaking colonies onto the same plates without uracil and leucine supplementation. After ∼48 h growth, yeast cells were collected and transferred into liquid medium containing YNB, CSM minus uracil and leucine, and 2% w/v raffinose for additional ∼48 hours. When OD_600_ reached ∼1.5, cells were diluted to OD_600_ around 0.5 into YPD media containing 2% w/v galactose for 12-16 h to induce DGAT1 overexpression. In each growth step in liquid media, cells were cultured at 30 °C at 250 rpm. To isolate DGAT1-containing microsomes, harvested yeast cells were resuspended in HSM buffer (50 mM HEPES pH7.5, 200 mM NaCl, 10 mM MgCl_2_) supplemented with protease inhibitor cocktail (Roche) and lysed by three passes to a prechilled microfluidizer at 1400 bar. Microsomal fractions were isolated by ultracentrifugation at 50,000g for 1h 4 °C. Membranes were resuspended in ice-cold HSM buffer supplemented with 10% v/v glycerol and snap frozen in liquid nitrogen for activity studies.

For crosslinking experiments, DGAT1 was expressed using previously established BacMam system^1^. Purified DGAT1 dimers in HSM buffer containing 0.05% w/v GDN was incubated with 5µM 1-palmitoyl-2-oleoyl-sn-glycerol (Avanti, 800815) or 5µM oleoyl coenzyme A (Sigma O1012) for 30 minutes at 37°C 500 rpm, followed by adding DSS crosslinker that freshly prepared in DMSO to a final concentration of 0.05 mM. After incubating for 30 min, the reaction was quenched by adding Tris-HCl, pH8.0 to a final concentration of 50 mM. The mixture was further analyzed by SDS-PAGE and western-blot analyses using DGAT1 primary antibody.

### DGAT1 activity assay

To access endogenous DAG bound with purified DGAT1, 2µg purified DGAT1 in GDN was supplemented with 50 µM oleoyl CoA containing 0.02 μCi [^14^C]-oleoyl-CoA as a tracer without adding exogenous DAG. As control, DGAT1 inhibitor A922500 was used at 10 µM. The activity assay of wild-type DGAT1 or mutants was performed according to our previous protocols^1,71^.

## Acknowledgments

We thank Julia Mahamid for supporting access to resources at EMBL and we thank EMBL Advanced Light Microscopy Facility for support in fluorescence image acquisition and analysis. S.V. acknowledges support by the Swiss National Science Foundation (PP00P3_194807, 310030_219264). This project has received funding from the European Research Council (ERC) under the European Union’s Horizon 2020 research and innovation program (Grant agreement No. 803952). This work was supported by grants from the Swiss National Supercomputing Centre (CSCS) under project ID s1011 and s1030. VTS was supported by the Marie Skłodowska-Curie Actions (01028297), the Biomedicum Helsinki Foundation, the Orion Research Foundation, and the Finnish Cultural Foundation. X.S. acknowledges support by the Robert Welch Foundation (grant A-2215-20240404).

All authors thank Valeria Zoni for many fruitful discussions and for a critical reading of the manuscript.

## Author contributions

Conceptualization, J.S., P.C., V.T.S., X.S. and S.V.; methodology, J.S., A.B., S.C.P., S.Y.W. and V.T.S.; Investigation, J.S., P.C., V.T.S., X.S. and S.V.; writing-original draft, J.S., P.C. and S.V.; writing-review & editing, J.S., P.C., V.T.S., X.S. and S.V.; funding acquisition, S.V., X.S. and V.T.S.; resources, S.V.; supervision, S.V.

## Supplementary material

**Supplementary Figure 1:**
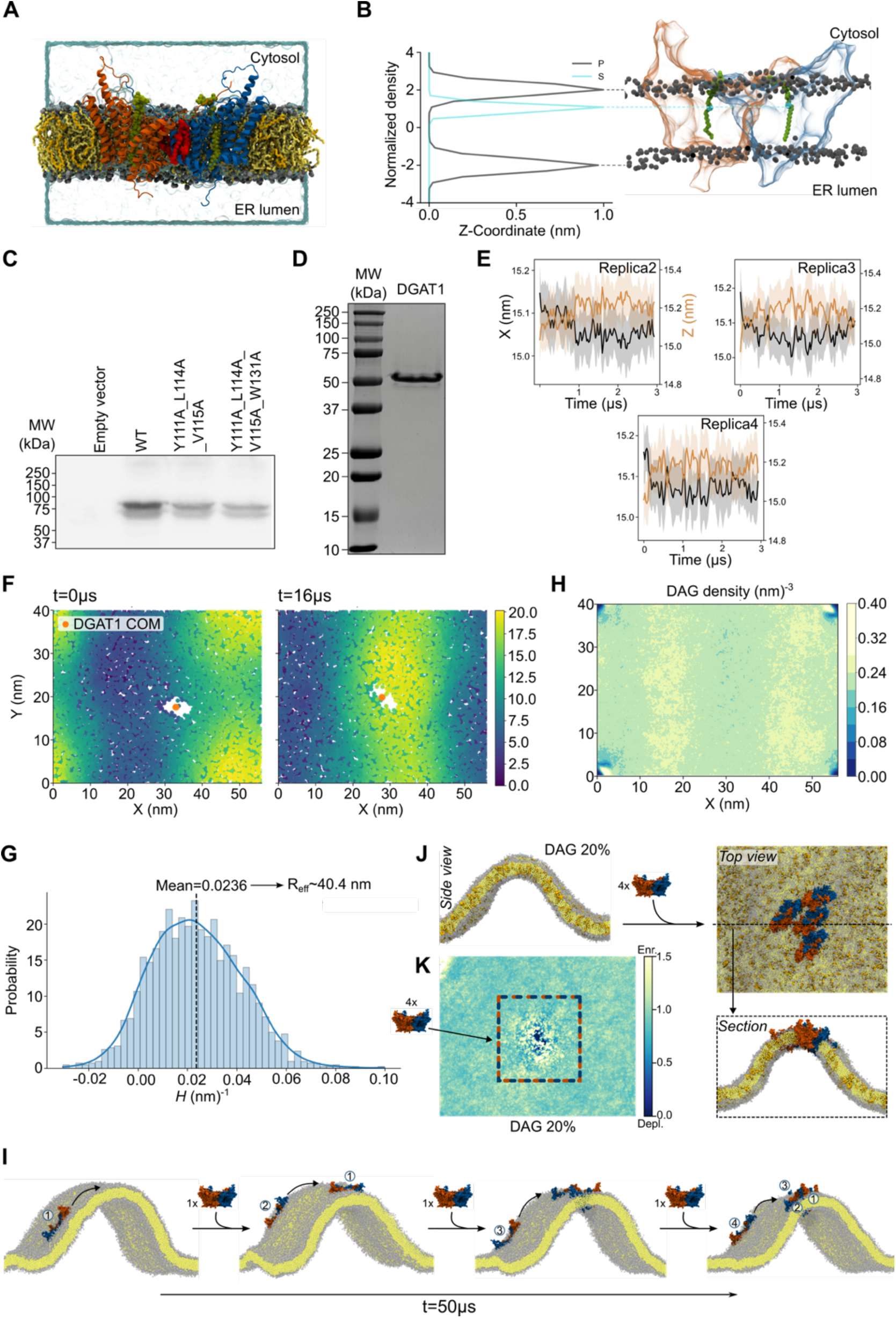
A) Representation of the solvated ER-like bilayer (DOPC: yellow; DAG: light orange) containing DGAT1 embedded bound to oleoyl-CoA (green) and POPE (red) molecules. **B)** The oleoyl-CoA molecules (green) position themselves at the level of the cytosolic leaflet, as clearly quantified by the localization of both oleoyl-CoA sulphur atoms (cyan) with respect to the phosphate groups (grey) of the lipid bilayer. **C)** Western blot analysis of yeast microsome overexpressing GFP-fused DGAT1 and mutants with anti-GFP antibody. **D).** SDS-PAGE analysis of purified DGAT1 in detergent. **E)** As in Fig. 3B, time evolution of the x (=y) and z box dimensions of systems composed by AA-DGAT1 embedded in flat DOPC membrane. **F)** As in Fig. 3D, DGAT1 homodimer’s (orange dot) propensity to localize in curved regions of a CG buckled bilayer. Analysis referred to a second independent replica. **G)** DGAT1 preferential localization in positively highly-curved regions in a CG buckled membrane showed as histogram of mean curvature (*H*). Values for mean curvature and effective radius are reported. **H)** Number density of DAG lipids showing their accumulations in curved regions of the buckled membrane (yellow areas) in a second independent MD production replica. **I)** Representation of the DGAT1 oligomerization workflow. The snapshots show the process through which DGAT1 dimer units have been embedded consecutively, one by one, into a CG buckled bilayer. In this exemple, the buckled membrane is constituted by 100% DOPC lipids, but the same workflow has been utilized for systems containing both 20% and 30% DAG lipids. **J)** Buckled membrane systems modelled in presence of DAG at 20% concentration, showing multiple copies of DGAT1 dimer clustering at the positively curved region. **K)** Enrichment map showing DAG accumulation in the proximity of the DGAT1 clusters 20% DAG concentration in a representative replica.

